# Superior colliculus peri-saccadic field potentials are dominated by a visual sensory preference for the upper visual field

**DOI:** 10.1101/2024.10.30.621170

**Authors:** Ziad M. Hafed

## Abstract

The primate superior colliculus (SC) plays important sensory, cognitive, and motor processing roles. Among its properties, the SC has clear visual field asymmetries: visual responses are stronger in the upper visual field representation, whereas saccade-related motor bursts are weaker. Here, I asked whether peri-saccadic SC network activity can still reflect the SC’s visual sensitivity asymmetry, thus supporting recent evidence of sensory-related signals embedded within the SC’s motor bursts. I analyzed collicular peri-saccadic local field potential (LFP) modulations and found them to be much stronger in the upper visual field, despite the weaker motor bursts. This effect persisted even with saccades towards a blank, suggesting an importance of visual field location. I also found that engaging working memory during saccade preparation differentially modulated the SC’s LFP’s, again with a dichotomous upper/lower visual field asymmetry. I conclude that the SC network possesses a clear sensory signal at the time of saccade generation.

## Introduction

The primate superior colliculus (SC) is critical for saccade generation^1–9^, but it also possesses a rich visual processing repertoire^10,11^. A now-acknowledged property of the SC is that it does not represent all visual field locations equally: besides foveal magnification^12,13^, SC neurons representing the upper visual field show significantly stronger and earlier visual responses than neurons representing the same retinotopic eccentricities in the lower visual field^14–16^. Intriguingly, saccade-related motor bursts in the same neurons exhibit the exact opposite asymmetry: SC motor bursts are stronger in the lower rather than upper visual field^14,17^. While this dichotomy between visual and motor responses suggests the presence of distinct SC neuronal processes associated with either visual sensation or saccade-related motor discharge^18–20^, other experiments have revealed evidence of sensory signals embedded directly within the motor bursts themselves^11,17,21–24^. In this study, I was particularly interested in further investigating such latter evidence.

Specifically, I asked whether a visual sensory preference for the upper visual field in the SC^14^ still manifests itself peri-saccadically, despite the weaker motor bursts. Since SC local field potentials (LFP’s) exhibit saccade-related modulations^21,25–27^, I investigated whether these modulations reflect the visual^14–16^ or the motor^14,17^ spiking asymmetries. As expected, I found that SC LFP’s exhibited strong peri-saccadic negativity at the time of motor bursts^21,25–27^. Critically, such negativity was much larger in the SC’s upper visual field representation than in its lower visual field representation, which directly matched the asymmetry in the SC’s stimulus-evoked LFP negativity. Peri-saccadic LFP modulations thus reflected the SC’s sensory preference for the upper visual field, and they were unambiguously dissociated from peri-saccadic spiking asymmetries.

I then checked whether this dissociation persisted even with saccades towards a blank, suggesting that topographic location within the SC could be a critical determinant for the peri-saccadic LFP modulation asymmetries. I employed memory-guided saccades, for which there was no visible stimulus either in the working memory delay period or, later, at the time of actual saccade generation. I found that SC LFP’s during the working memory delay period still exhibited a clear upper/lower visual field asymmetry. Importantly, peri-saccadic LFP negativity in the absence of a visible saccade target was still significantly stronger for the SC’s upper, rather than lower, visual field representation.

My results suggest that at the time of saccades, the local SC network possesses a clear sensory-related signal, potentially relevant for processes beyond just online saccade control.

## Results

### Stronger peri-saccadic field potential deflections in the upper visual field

We previously documented an asymmetry in the SC’s representation of the upper and lower visual fields^14^. Here, I revisited the same database^14^, but now focusing on peri-saccadic LFP modulations. To better orient the readers, I first summarize how the stimulus-evoked LFP’s behaved. Figure 1A reproduces earlier results^14^, demonstrating that visually-evoked LFP modulations were stronger (more negative) in the upper visual field. Note that in each experiment, the appearing stimulus was always placed at the best response field (RF) location of the recorded neurons^14^, and that single-electrode penetrations were performed in these experiments (Methods). The inset, again reproduced from ref. ^14^, shows that this upper/lower visual field LFP asymmetry was consistent with the asymmetry in SC visual response firing rates: visual bursts were stronger in the upper visual field, just like LFP negativity was larger. Figures 1B, C, in turn, remind that the LFP asymmetry was step-like across the horizontal meridian^14^, suggesting a potential quadrant-based discontinuity in the SC’s topographic representation of the retinotopic visual field.

**Figure 1.**
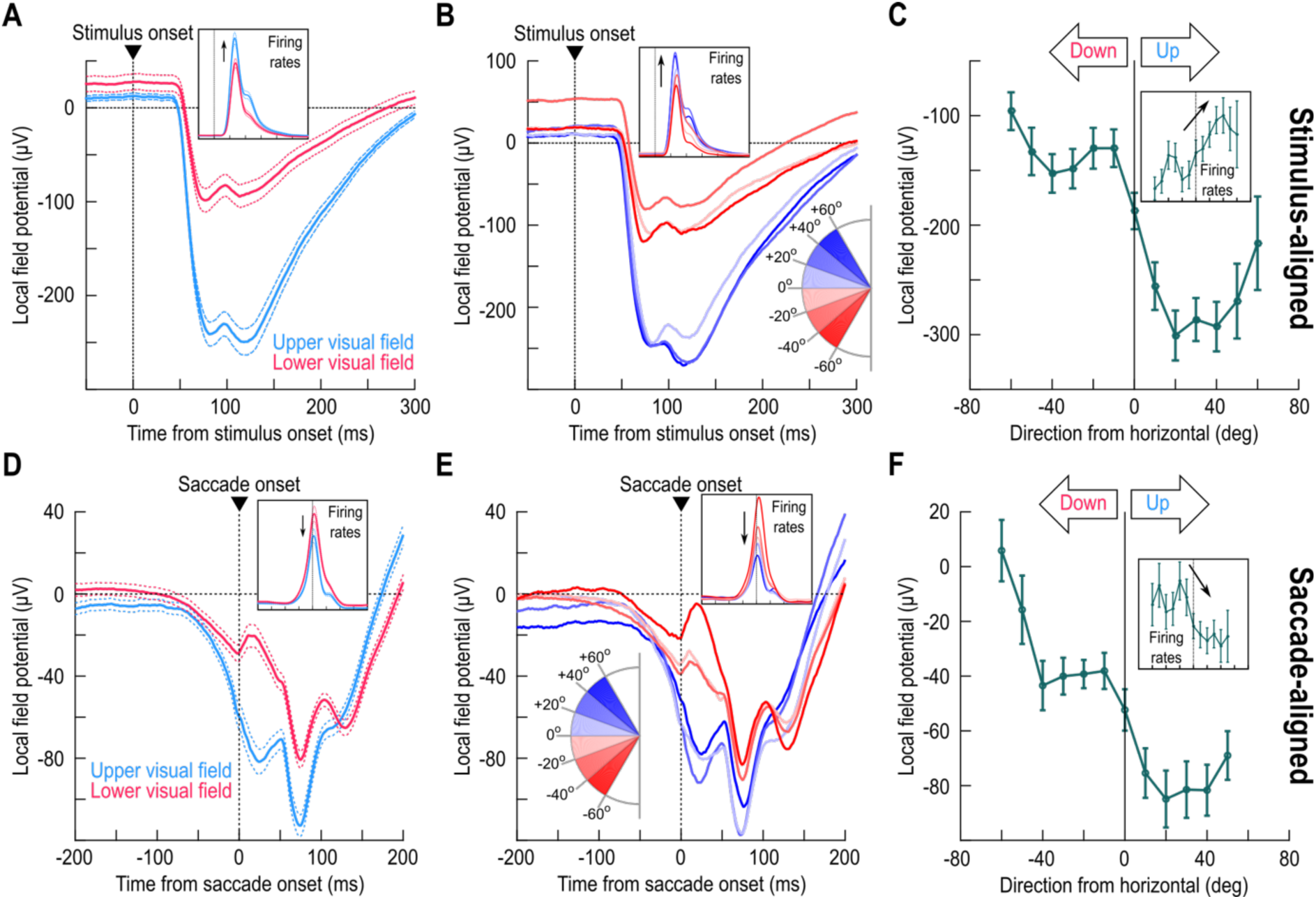
Peri-saccadic local field potential (LFP) deflections in the superior colliculus (SC) reflect their visual counterparts. **(A)** Stimulus-evoked LFP deflections after the brief appearance of a small white spot at the preferred response field (RF) location^14^. Blue shows the average for all electrode tracks in the SC’s upper visual field representation; red shows results from the lower visual field representation. The inset shows the corresponding single-neuron firing rates^14^. LFP negativity deflections were much stronger in the upper visual field representation^14^. **(B, C)** The same data but now separated as a function of direction from the horizontal axis represented by the SC sites. There was a step-like discontinuity across the horizon^14^. **(D)** Similar to **A** but now for data aligned to saccade onset towards a visible spot. Even though the motor bursts were weaker in the upper visual field^14,17^ (inset), the LFP negativity deflections were still much stronger for the upper visual field, consistent with the stimulus-evoked effects in **A**. **(E, F)** The same discontinuity across the horizontal meridian seen in **B**, **C** was also evident peri-saccadically (0-50 ms from saccade onset in **F**; Methods). Note how the insets in **B**, **E** as well as those in **C**, **F** show directly opposing dependencies in the spiking activity between the visual and motor epochs; in contrast, the LFP’s show the same dependencies. Error bars in all panels denote SEM. **A**-**C** were reproduced (using updated color schemes) with permission from our earlier study^14^ for easier direct visualization next to **D**-**F**. All insets were also reproduced from the same study^14^.

Peri-saccadically, we previously noted that SC motor bursts are actually weaker in the upper visual field^14,17^, an opposite asymmetry from the visual bursts. Does that mean that LFP negativity would also be smaller? Using a delayed, visually-guided saccade task (Methods), I plotted in Fig. 1D average peri-saccadic LFP modulations across all tested electrode tracks. Like in the stimulus-evoked analyses (Fig. 1A), I classified electrode tracks according to whether they penetrated the SC’s upper (blue) or lower (red) visual field representation. Also like in the stimulus-evoked analyses, the saccade target was always at the best RF location of the session^14^. There was a much larger peri-saccadic LFP negativity in the upper visual field representation (p=1.6612×10^-9^; t-test for the interval 0-50 ms from saccade onset, comparing upper and lower visual field electrode tracks). This was despite the weaker motor bursts (inset). Thus, peri-saccadically, aggregate SC network activity around the recording electrodes reflected the strong visual sensory preference for the upper visual field, and not the weaker simultaneously-occurring motor bursts – a clear dissociation between SC spiking and LFP characteristics.

Given this dissociation, and given that LFP signals might reflect the influence of synaptic and other network activity^28–35^, these results suggest that SC peri-saccadic sensory information could act to weaken motor bursts. Interestingly, we recently found a potential inverse relationship between image preference at stimulus versus saccade onset in visual-motor SC neurons: for some image manipulations, if a neuron fired more for one image feature in its visual burst, it tended to also have weaker saccade bursts for the same feature^21^.

Remarkably, the visual field asymmetry in peri-saccadic LFP negativity also exhibited a step-like discontinuity across the horizon: just like in the stimulus-evoked LFP’s (Fig. 1B), all tested electrode tracks in the upper visual field representation had similar peri-saccadic LFP negativity amplitude, which was larger (in absolute value) than all tested electrode tracks in the lower visual field representation (Fig. 1E). This can also be seen in Fig. 1F; here, I measured the LFP amplitude in the interval 0-50 ms from saccade onset. There was a similar dependence to the stimulus evoked modulations in Fig. 1B, C (p=3.392×10^-12^; one-way ANOVA across the shown angular direction bins of Fig. 1F). Again, this dependence was the opposite of that in the spiking activity (inset). And, this dependence persisted even in the immediate pre-saccadic^3,26,36^ interval (Fig. S1).

It is also worthwhile to consider the magnitude of the peri-saccadic LFP asymmetry. From Fig. 1D, F, I calculated the LFP amplitude in the interval 0-50 ms from saccade onset; it was ∼2.7 times larger (in absolute value) in the upper, rather than lower, visual field representation of the SC. This was similar to the stimulus-evoked LFP asymmetry of Fig. 1A-C (peak LFP amplitude in the interval 30-100 ms from stimulus onset being ∼2.5 times larger in the upper visual field^14^). Thus, despite the quantitative and qualitative differences in visual field asymmetries for stimulus-evoked versus saccade-related spiking, the LFP asymmetries were both qualitatively and quantitatively similar.

The peri-saccadic LFP asymmetry was also present in all ranges of tested eccentricities and directions. For example, just like we did earlier for stimulus-evoked effects^14^, I measured LFP amplitude in the interval 0-50 ms from saccade onset and plotted it as a function of either eccentricity or direction bins represented by the electrode tracks. I also did the same analyses, which were not documented in the earlier study^14^, for the motor burst spiking. In every eccentricity (Fig. 2A) and direction (Fig. 2B) bin, motor bursts were weaker in the upper visual field^14,17^ (p=0.0151 and p=0.2659 for main effects of upper/lower visual field and eccentricity, respectively, in Fig. 2A; p=0.0001 and p=0.2319 for main effects of upper/lower visual field and direction from horizontal, respectively, in Fig. 2B; two-way ANOVA in each case; Methods). This effect was reversed for LFP’s, and also amplified (Fig. 2C, D) (p=0 and p=0.7458 for main effects of upper/lower visual field and eccentricity, respectively, in Fig. 2C; p=0 and p=0.2191 for main effects of upper/lower visual field and direction from horizontal, respectively, in Fig. 2D; two-way ANOVA in each case). Using the same analysis approach, I also checked the LFP and spiking modulations during the immediate pre-saccadic interval^3,26,36^: once again, the LFP’s, but not the spikes, still reflected the SC’s sensory preference for the upper visual field (Fig. S2).

**Figure 2.**
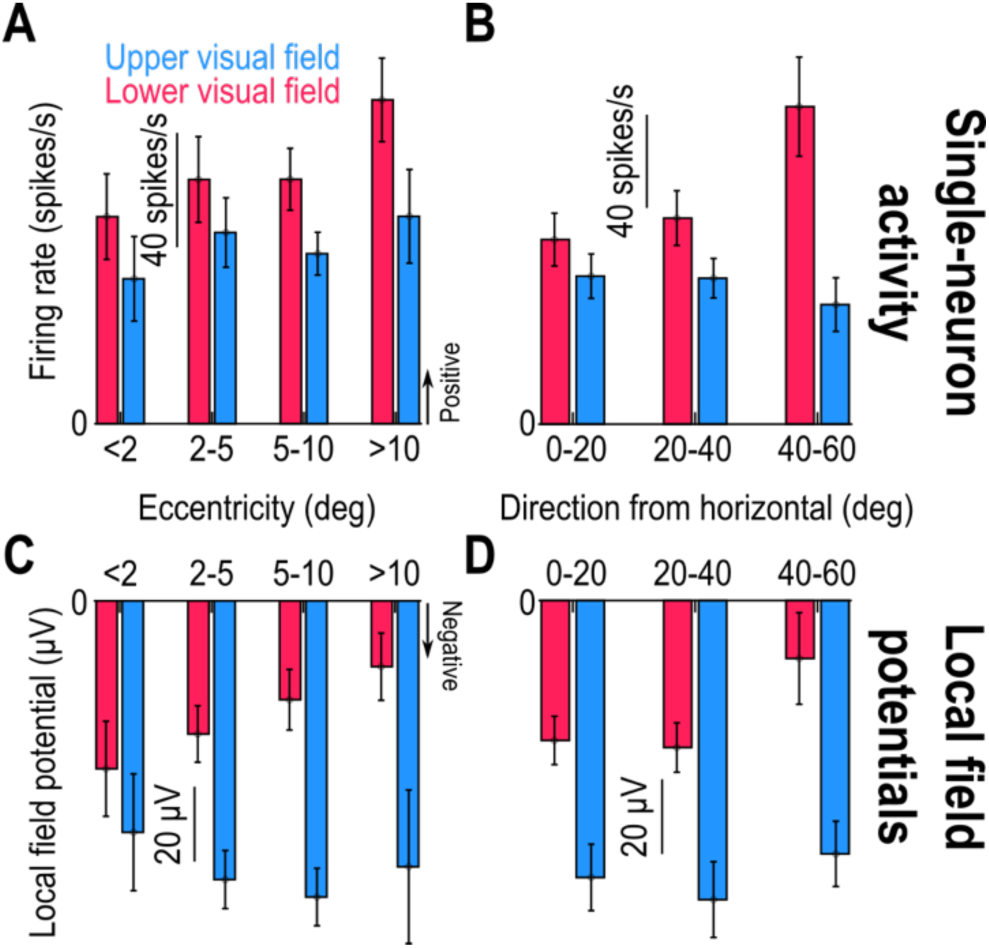
Consistency of the results of Fig. 1 across tested eccentricities and directions. **(A, B)** For all single neurons from the database, I binned them according to the eccentricity (**A**) or direction from horizontal (**B**) of their RF hotspots (similar binning to what we had done with visually-evoked LFP measurements in our earlier study^14^). This confirmed the consistently weaker motor bursts in the upper visual field^17^. In each bin, the measurements were made in the interval 0-50 ms from saccade onset. **(C, D)** For peri-saccadic LFP’s, there was always a stronger negativity for the upper visual field, like in the stimulus-evoked deflections (Fig. 1) and opposite to the spiking asymmetry of **A**, **B**. Error bars denote SEM. Figures S1, S2 document similar results when measuring pre-saccadic activity.

Thus, I observed a strong asymmetry in the SC’s peri-saccadic LFP’s, and this asymmetry was the same as that in both the spiking and LFP sensory-driven responses (Fig. 1)^14^; this asymmetry was opposite of that in the spiking motor bursts of the SC.

### Visual field location as a defining feature

In light of our recent observations that there is a sensory representation embedded within both SC motor bursts and peri-saccadic LFP’s^21^, the results above imply that stimulus location may itself be considered a visual feature. If so, then it is the SC neurons’ topographic location, and not stimulus presence per se, that should matter. To confirm this, I analyzed data from memory-guided saccades with no visible saccade target (Methods). I first investigated how invoking working memory^37^ per se affected the delay-period LFP values (well before saccade generation), and I then focused on the peri-saccadic LFP negativity modulations that I was primarily interested in. In both cases, it became clear that the SC topographic location did indeed matter, as I predicted.

Engaging working memory strongly altered the SC’s LFP patterns during the delay period leading up to saccade generation^25^, but, remarkably, it still did not eliminate the SC’s upper/lower visual field dichotomy. Consider, for example, Fig. 3A, B. In Fig. 3A, I aligned the LFP data to the end of the delay period, when the “go” signal for the saccade was provided (Methods), and I did this for the visual condition (with a visible saccade target throughout the whole trial). In Fig. 3B, I did the same, but now for the memory-guided saccade condition (Methods), in which there was no visible saccade target during the delay period. Consistent with earlier reports^25^, the memory condition caused a substantial upward shift in pre-saccadic LFP levels when compared to the visual condition (compare the corresponding curves in Fig. 3A, B). Interestingly, the delay-period LFP’s in both cases were still dependent on the visual field location represented by the recorded SC sites. For example, in the visual condition, LFP’s were more negative in the upper visual field SC sites than in the lower visual field SC sites (Fig. 3A). This is the same dependence as that observed in Fig. 1A, suggesting a sustained influence of continuous visual stimulation on the SC’s LFP’s. In the case of memory-guided saccade preparation, LFP amplitudes (positive this time^25^) were higher (more positive) for the upper visual field SC sites than for the lower visual field sites (Fig. 3B). Thus, delay-period SC LFP’s depended on visual field location in both the visual and memory conditions, but the sign of the dependence was reversed in the memory condition.

**Figure 3.**
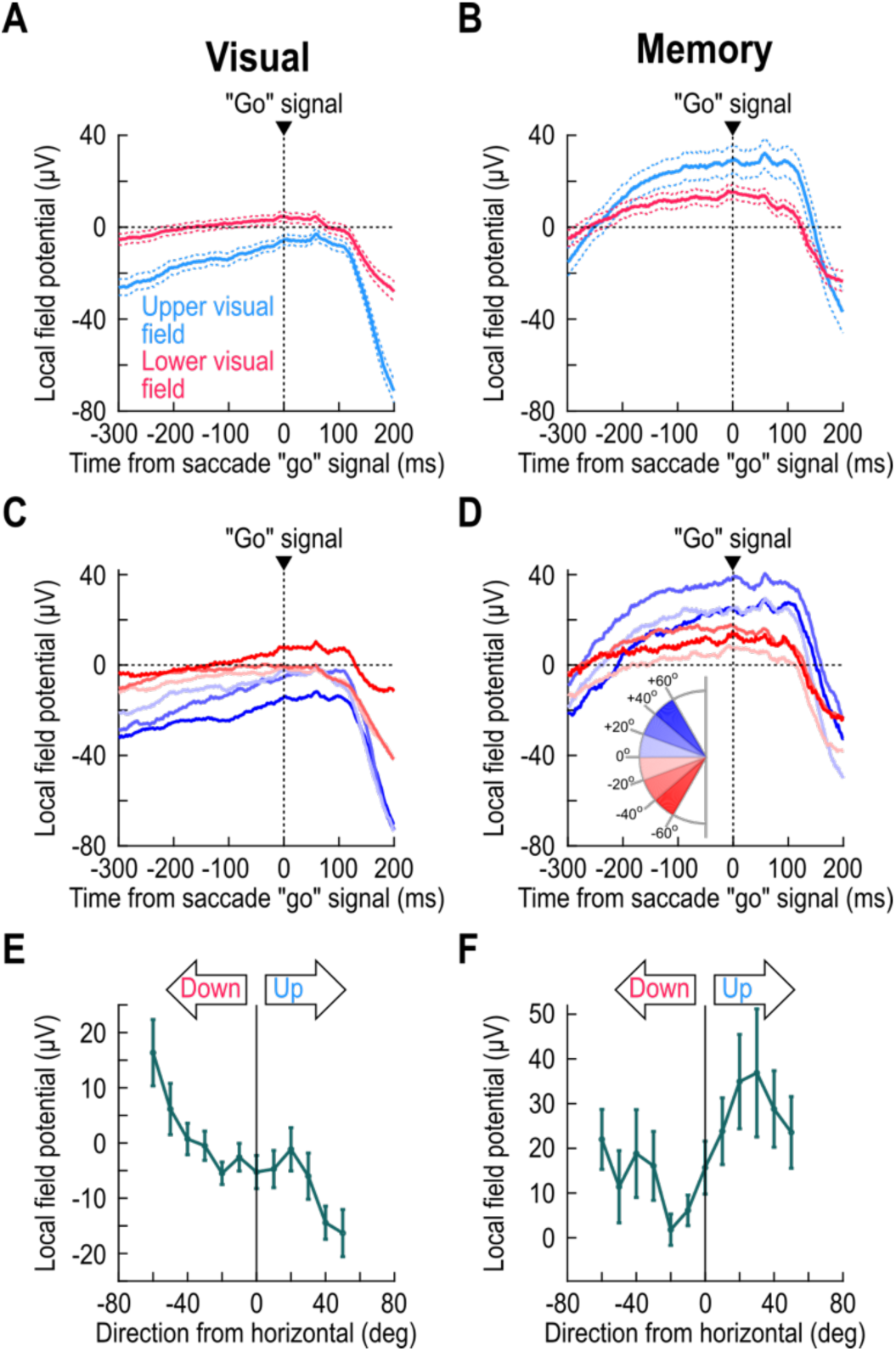
Both visual and working memory LFP representations depend on visual field location, but in diametrically opposite ways. **(A)** Same as Fig. 1D but aligned to the end of the delay period that existed between stimulus onset and the “go” signal for the saccade (in the delayed, visually-guided saccade task; Methods). Here, I aligned the data to the time of the go signal, when the fixation spot was removed and the eccentric spot (saccade target) was still visible. Before the end of the delay period, there was a sustained effect in which there was stronger LFP negativity for upper visual field SC sites. This is consistent with the results of Fig. 1A-C and suggests that continuous visual stimulation has persistent effects on the SC’s LFP’s. The strong negative deflections ∼150-200 ms after the go signal reflect the peri-saccadic LFP modulations of Fig. 1D-F. **(B)** In the absence of a visible saccade target, LFP’s turned positive after the initial negativity transient associated with the earlier stimulus onsets; thus, the SC’s LFP’s were clearly positive by the end of the delay period^25^. Interestingly, there was still a dependence on the visual field location: SC sites representing the upper visual field had a larger LFP positivity during working memory than SC sites representing the lower visual field. Figure 4 shows that, despite this difference in LFP signals at the time of the go signal for memory-guided saccades, later peri-saccadic negativity amplitudes still behaved like those in the visually-guided saccade task. **(C, D)** Same as **A**, **B**, but now separating electrode tracks as a function of the direction from horizontal represented by the targeted SC sites. For the visual condition (**C**), the same sensory dependence of Fig. 1A-C was observed, again suggesting a sustained sensory influence on SC LFP’s. In the case of memory (**D**), there was clearly a visual field effect, but this time of the opposite sign: upper visual field SC sites had stronger LFP positivity, rather than negativity, at the end of the delay period. **(E, F)** Same as Fig. 1C, F, but this time when measuring LFP values at the end of the delay period in both tasks (from -50 ms to +25 ms relative to the time of the saccade go signal; Methods). In both cases, there was an upper/lower visual field effect, but having different signs. Error bars denote SEM. Also see Fig. S3.

I confirmed the above observations statistically. In the visual case (Fig. 3A), I gathered all electrode tracks targeting the SC’s upper visual field representation into one group, and I compared their delay-period LFP levels (averaged over the interval from -50 ms to +25 ms relative to the saccade go signal; Methods) to those from the electrode tracks targeting the lower visual field representation. Upper visual field LFP levels were significantly more negative than for the SC’s lower visual field representation (p=4.235×10^-4^; t-test; Fig. 3A), again implying a sustained influence of continuous visual stimulation on SC LFP’s. For the memory case (Fig. 3B), there was still a significant difference between the upper and lower visual field SC sites (p=0.0204; t-test); this time, however, the upper visual field electrode tracks had significantly more positive, rather than more negative, LFP levels (Fig. 3B).

I also separated electrode tracks based on the direction from horizontal that their SC sites represented, just like I did in Figs. 1, 2 above. There was still evidence of a functional discontinuity^14^ across the horizontal meridian in both the visual (Figs. 3C, E, S3A) and memory (Figs. 3D, F, S3B) conditions, with the only difference being that, in the memory condition, upper visual field SC sites were associated with more LFP positivity, rather than more LFP negativity, when compared to lower visual field SC sites (p=6.913×10^-5^ and p=0.0236 for the visual and memory conditions, respectively; in each case, one-way ANOVA across the shown angular direction bins of either Fig. 3E or Fig. 3F). The pressing question, then, was whether, in the memory condition, these effects eliminated, or otherwise altered, the stronger peri-saccadic negativity that I observed so clearly in the SC’s upper visual field representation with visually-guided saccades (Figs. 1-2).

Peri-saccadically, the difference in LFP negativity between the upper and lower visual fields still persisted even in the absence of a visible saccade target. Figure 4A, B shows peri-saccadic LFP’s for both visually-guided and memory-guided saccades. In the upper visual field, the memory condition caused a substantial shift in pre-saccadic LFP voltage, as expected from Figs. 3A, C, E, S3A. This shift was essentially absent in the lower visual field (Figs. 3B, D, F, S3B), corroborating the idea of substantive functional differences between the SC’s upper and lower visual field representations^14,15^ (Fig. 3). More importantly, peri-saccadically, the amplitude of the deflection from pre-saccadic baseline levels was bigger in the upper than lower visual field, consistent with Figs. 1, 2. This observation is better appreciated after applying a baseline shift (Fig. 4C; Methods): there was still a clearly stronger negativity for upper visual field sites (p=0.003; t-test for the interval 0-50 ms from saccade onset, comparing upper and lower visual field electrode tracks), which was also present in different direction bins (Fig. 4D). Indeed, at all tested eccentricity (Fig. 4E) and direction (Figs. 4F, 5) bins, there was stronger peri-saccadic LFP negativity for the upper visual field (p=0.0048 and p=0.5844 for main effects of upper/lower visual field and eccentricity, respectively, in Fig. 4E; p=0.0188 and p=0.3152 for main effects of upper/lower visual field and direction from horizontal, respectively, in Fig. 4F; two-way ANOVA in each case). Thus, SC topographic location was the critical feature, even in the absence of a visible saccade target.

**Figure 4.**
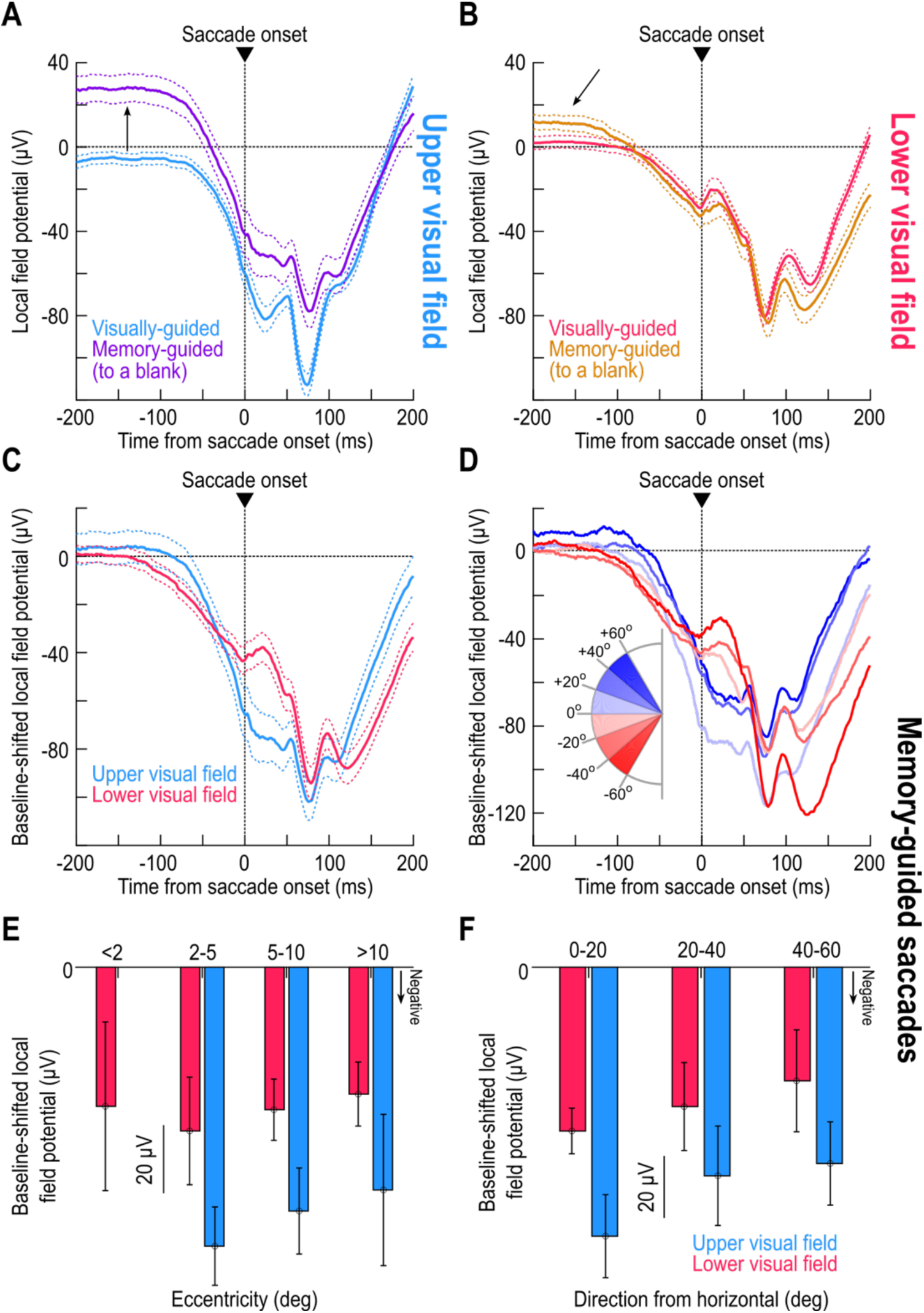
Stronger upper visual field peri-saccadic LFP modulations even in the absence of a visible saccade target. **(A)** Peri-saccadic LFP’s for upper visual field electrode tracks with and without a visible target (the visible target curve is the same as that in Fig. 1D). Long before the saccade (upward arrow), the memory condition elevated LFP voltages^25^ (Fig. 3). However, the peri-saccadic deflection magnitude was similar in the two curves. **(B)** Same as **A** but for the lower visual field. Here, the memory condition had a much weaker effect long before saccade onset (oblique arrow) (Fig. 3). Importantly, the peri-saccadic deflection strength was still similar in both conditions. **(C)** Applying a baseline shift in the memory condition (Methods) revealed a similar peri-saccadic asymmetry between the upper and lower visual fields in the absence of a visible saccade target; there was stronger negativity in the upper visual field. **(D)** Same as Fig. 1E but for the memory condition (after baseline shift). Upper visual field tracks had consistently stronger peri-saccadic LFP negativity. For example, compare the sites directly straddling the horizontal meridian (least saturated blue and least saturated red), especially in the peri-saccadic interval. **(E, F)** Same as Fig. 2C, D, but for the memory condition, again showing stronger negativity in the upper visual field (the missing bar for <2 deg eccentricity means no data). Error bars in all panels indicate SEM.

**Figure 5.**
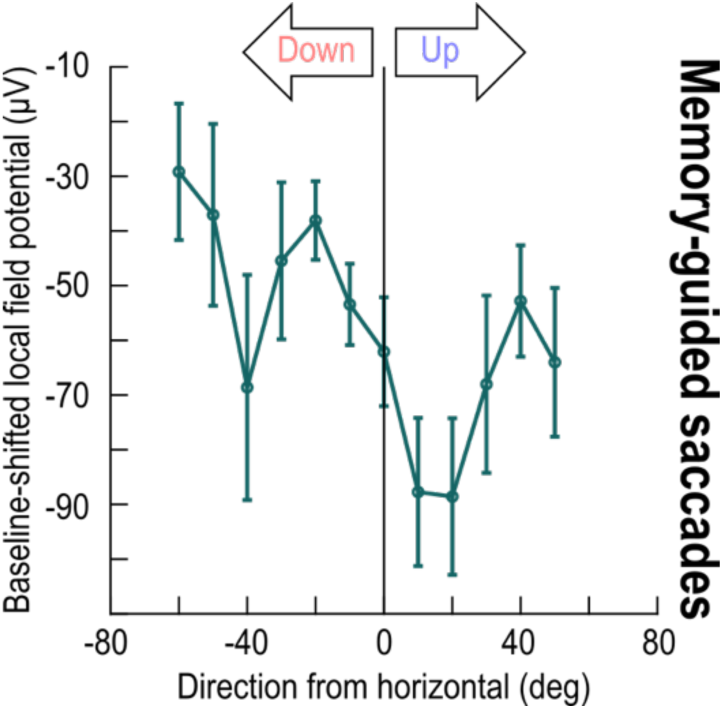
Direction-dependence of peri-saccadic LFP modulations in the absence of a visible saccade target. This figure bins all electrode tracks from Fig. 4 as a function of the direction from horizontal represented by the recorded SC neurons. For baseline-shifted peri-saccadic LFP measurements (during the interval of 0-50 ms from memory-guided saccade onset), there was a stronger LFP negativity in the upper visual field representation of the SC than in the lower visual field representation. This was confirmed statistically when I grouped all upper visual field electrode tracks into one group and all lower visual field electrode tracks into another (p=0.003; t-test). Similarly, a one-way ANOVA across the shown angular direction bins revealed a significant effect of direction from the horizontal meridian (p=0.0123). Error bars denote SEM.

### Upper visual field saccades to a blank are still associated with weaker motor bursts

Armed with the results of Figs. 1-5, I could now predict that SC motor bursts should be weaker in the upper visual field even for memory-guided saccades. I confirmed this: I consistently observed weaker saccade-related motor bursts for upper visual field SC neurons (Fig. 6) (p=0.0111 and p=0.202 for main effects of upper/lower visual field and eccentricity, respectively, in Fig. 6C; p=0.0007 and p=0.2876 for main effects of upper/lower visual field and direction from horizontal, respectively, in Fig. 6D; two-way ANOVA in each case).

**Figure 6.**
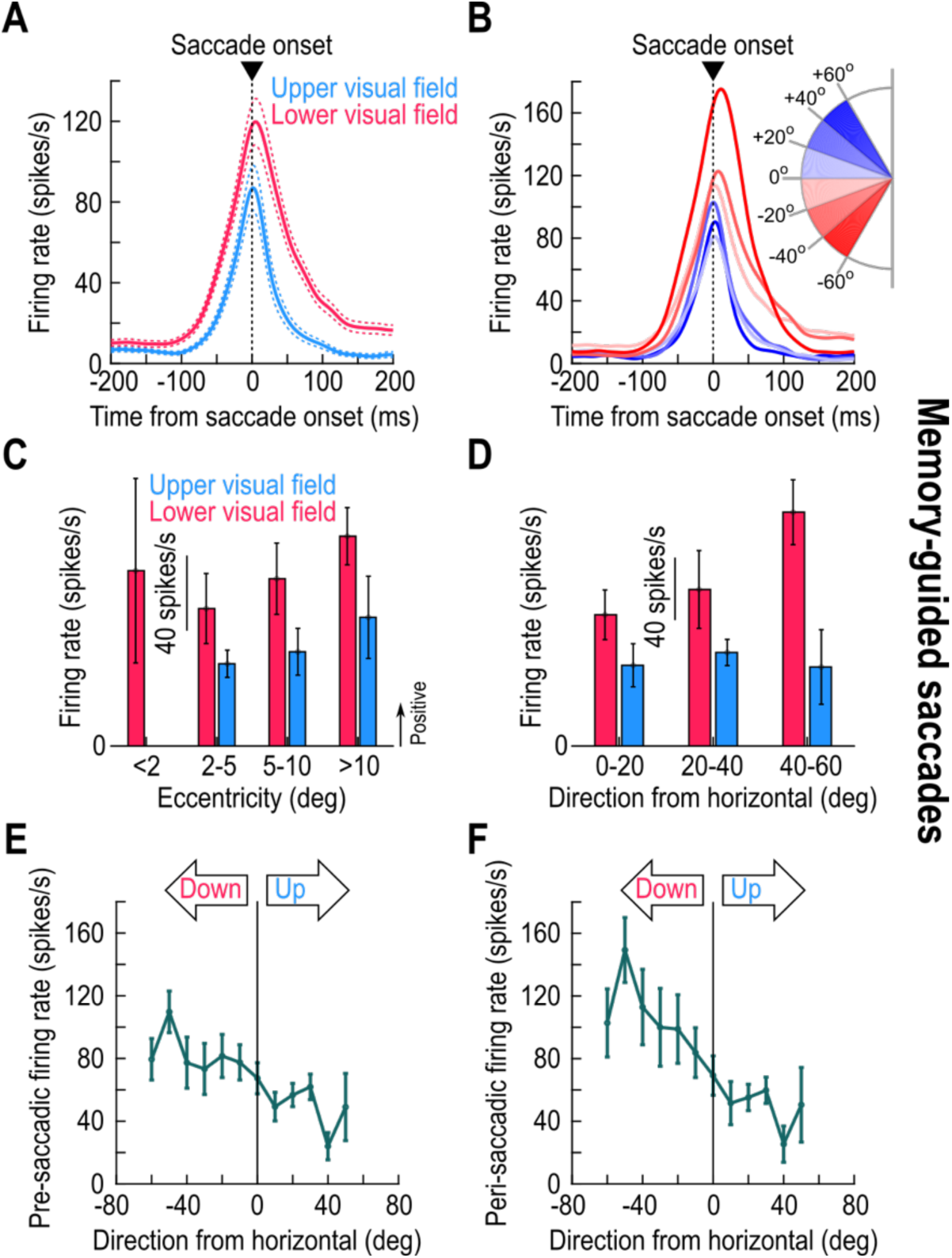
Weaker motor bursts in the upper visual field in the absence of a visible saccade target. **(A,B)** Like the insets of Fig. 1D, E but for the memory-guided saccade task. Note how the pre-saccadic firing rates did not depend on visual field location, even though the pre-saccadic LFP effects did (Figs. 3, 4A, B). This suggests another dissociation between SC LFP’s and spiking activity, beyond the dissociation I highlighted above for peri-saccadic intervals (e.g. Figs. 1, 2, S1, S2). **(C, D)** Like Fig. 2A, B but for memory-guided saccades. Motor bursts in the upper visual field were weaker than in the lower visual field, which is opposite the peri-saccadic LFP asymmetries (Figs. 4, 5). **(C)** Similar to Figs. 1C, F, 3E, F, 5, S1, except that I now measured firing rates in the final 50 ms before saccade onset. Pre-saccadic firing rates were weaker in the upper than in the lower SC visual field representation, and this happened even with saccades towards a blank (memory-guided saccades). I confirmed this with a t-test comparing all upper visual field neurons in the database in one group to all lower visual field neurons in the other (p=0.0039). I also performed a one-way ANOVA across all shown angular direction bins, again with a significant effect (p=0.0329). **(D)** Same as **C** but now for the measurement interval 0-50 ms from saccade onset. Again, peri-saccadic firing rates were weaker in the upper visual field for memory-guided saccades (p=0.0037; one-way ANOVA across all shown angular direction bins). Note that the overall firing rates were generally higher than in **C** (same y-axis ranges in the two panels), which is expected because motor bursts are known to reach peak firing rates some time peri-saccadically rather than pre-saccadically^3,36^. Error bars denote SEM.

Interestingly, the pre-saccadic spiking activity of memory-guided saccades did not depend on the visual field location (Fig. 6A, B) unlike in the delay-period LFP levels (Fig. 3B, D, F). This observation was also true for the visually-guided saccades (as depicted in our earlier publications^14,17^). Moreover, in Fig. 4A, B of ref. ^14^, it was clear that visual spiking activity during the memory epoch had subsided to similar levels for either upper or lower visual field SC neurons, even though delay-period LFP amplitudes were still differentiated (Figs. 3A, C, E, S3A). Thus, combined with Fig. 3 above, and across both the visual and memory conditions, I observed yet another interesting dissociation between delay-period SC spiking and LFP dependencies, beyond the peri-saccadic dissociation of Figs. 1, 2, 4.

In summary, my results suggest that peri-saccadic local SC network activity: (1) reflects the strong sensory preference for the upper visual field (Figs. 1-3); (2) is accompanied by weaker spiking bursts (Figs. 1, 2, 6); and (3) is a function of SC topography (Figs. 3-5). Thus, sensory tuning in SC movement commands^21^ extends to the domain of visual field asymmetries.

## Discussion

If SC peri-saccadic LFP modulations directly reflect spiking activity in saccade-related motor bursts, as might be generally assumed^25^, then I should have observed weaker peri-saccadic negativity for the upper visual field SC representation, concomitant with weaker motor bursts^14,17^. I found the opposite: peri-saccadic LFP modulations reflected the SC’s sensory, rather than motor, preference. I believe that these results add to our recent evidence^21^ of sensory tuning in SC motor bursts.

The fact that LFP’s can be dissociated from spiking activity has been observed before, even in the SC itself, albeit in the context of signal timing. Specifically, in stimulus-evoked responses, LFP modulations normally lead stimulus-driven spiking in both the SC^26^ and frontal eye fields (FEF)^38^, suggesting that the spiking activity might reflect the local network’s processing of incoming sensory signals. On the other hand, peri-saccadically, LFP modulations lag the onset of spiking motor bursts^26,38,39^. Thus, the timing relationship between LFP’s and spiking is flipped peri-saccadically. In my case, I found an even more remarkable dissociation: peri-saccadic LFP modulations reflected the SC’s stimulus-evoked preferences (whether in LFP’s or spiking) and not the motor burst preferences. Thus, SC sensory preferences for the upper retinotopic visual field seem omnipresent in LFP’s.

I also observed a secondary dissociation between SC spiking activity and LFP’s in my analyses. Specifically, well before saccade onset, firing rates were similar across different visual field locations represented by the recorded SC sites, both for memory-guided saccades (Fig. 6B) as well as for visually-guided saccades (refs. ^14,17^). However, the LFP levels at these pre-saccadic times (near the end of the delay period) were clearly dependent on visual field location (Figs. 3, 4A, B). These differences in LFP’s as a function of visual field location could represent a latent signal, which may not be overtly visible in the spiking, but which can be “uncovered” under specific behavioral scenarios. For example, using a trigeminal blink reflex to trigger premature eye movements during delay periods, Jagadisan and Gandhi^40^ discovered that classic “motor neurons” of the SC possessed a hidden visual signal present within them. It would be interesting to try to experimentally disrupt the working memory phase of memory-guided saccade paradigms (for example, by using strategically timed visual stimulation^41–43^) to try to discover a functional significance for the latent LFP effects that I observed in Fig. 3.

Returning back to the peri-saccadic LFP’s, which represent the main topic of my study, I interpret my results as being consistent with our recent observations of a sensory signal embedded within SC motor bursts^21^. In line with this, we do know that SC motor bursts vary as a function of visual field location^14,17^. However, there is one difference that is worth noting here: while SC visual responses are stronger for upper visual field stimuli^14^, saccade-related motor bursts are weaker^14,17^. This suggests a model in which SC network activity is actively reformatted^18,19^ between the visual and motor epochs, and I believe that such reformatting could happen for a variety of reasons beyond saccade control. For example, one kind of reformatting could help downstream areas to correctly distinguish between SC visual and motor bursts, which is important in order to avoid unwanted saccades at the time of “visual” spikes^20,21,44,45^. My results add another distinct possibility: the reformatting could be relevant for corollary discharge and its influences on peri-saccadic vision. That is, if there is a sensory signal present in the SC peri-saccadically^21^, and if this signal is not relevant for controlling the eye movements themselves^17^, then this signal could be part of what is relayed^46–49^ from the SC to other brain areas. In turn, this signal could alter peri-saccadic vision in strategic ways^15,50^. Interestingly, we recently found that peri-saccadic visual sensitivity becomes momentarily better in the upper visual field, which is opposite from how it behaves without saccades^15^. Thus, a peri-saccadic sensory signal in the SC could be used to transitorily alter well-known visual field perceptual asymmetries^51–53^.

I am also intrigued by the fact that I observed the same peri-saccadic effects with memory-guided saccades. This suggests that topographic location in the SC’s functional map of the visual field is what matters, and it is also consistent with the idea that visual location is itself an important “feature” for vision. This observation is also in line with the idea that LFP’s might reflect local anatomy and connectivity^28–32,34,35^, especially if one were to consider that the SC might amplify its anatomical representation of the upper visual field^14^. Indeed, for recording sites away from the eye movement endpoints, peri-saccadic LFP modulations are much smaller in amplitude^54^ than what I observed here.

Finally, my analyses of memory-guided saccades revealed an interesting property of delay-period LFP modulations in the SC. It was previously known that invoking working memory during the memory-guided saccade paradigm causes an elevation of SC LFP levels right after the transient, sensory-evoked LFP negativity^25^. Not only did I replicate these observations here (Fig. 3B, D), but I also found that this LFP hallmark of working memory allocation in the SC is strongly dependent on visual field location (Figs. 3D, F, S3B). I believe that this is an interesting discovery, especially because it can help clarify some previously documented idiosyncrasies of memory-guided oculomotor behavior. For example, memory-guided saccades tend to always have a systematic bias to land slightly above the true remembered target locations^55–57^. Similarly, eye position and other oculomotor properties can be biased upward under darkness conditions^58,59^. It would be interesting in future studies to relate my observations about upper/lower visual field asymmetries in delay-period LFP’s to these oculomotor phenomena. More importantly, it would be interesting to explore whether such SC asymmetries can explain biases and asymmetries in working memory paradigms that do not explicitly require an overt oculomotor response. If biases emerge in these paradigms that are congruent with the SC LFP biases that I found, then this can help support the idea that the SC may be considered to be a driver for cognitive processes, like covert visual attention.

In all, my results motivate further studies of peri-saccadic visual field asymmetries, both perceptually and at the neuronal level.

## Limitations of the study

In this study, I only had access to single-electrode recordings (Methods). Therefore, I could not simultaneously record LFP’s across different SC layers. Such simultaneous recordings could allow performing current source density (CSD) analyses^27,60,61^, which could in turn help clarify exactly what electrophysiological processes near electrode contacts are captured by the SC LFP’s. For example, we can potentially discover a plausible interpretation for the dissociations between spiking and LFP asymmetries that I observed in my current study. I believe that this is a worthwhile future direction. For example, using simultaneous depth recordings, we recently did find that different SC functional layers had slightly different patterns of peri-saccadic LFP deflections^21^, suggesting that CSD analyses would indeed reveal candidate sources and sinks across the depth of the SC.

It might also be worthwhile in the future to consider spontaneous saccades, or saccades in complete darkness. This can allow further dissociation between motor bursts and sensory stimulation, and it can also allow avoiding the relatively unnatural delay period that is employed in classic laboratory visually-guided and memory-guided saccade tasks. On the other hand, spontaneous saccades forego experimental control over instantaneous cognitive state, requiring more careful planning of data analyses.

## Resource availability

### Lead contact

Further information and requests for resources and reagents should be directed to and will be fulfilled by the lead contact, Ziad Hafed.

### Materials availability

This study did not generate new unique reagents.

### Data and code availability

All data reported in this paper will be shared by the lead contact upon request.

This paper does not report original code.

Any additional information required to reanalyze the data reported in this paper is available from the lead contact upon request.

## Acknowledgements

I was funded by the Medical Faculty of the University of Tübingen.

## Author contributions

Z.M.H. designed the experiments and wrote the paper.

## Supplemental information

Document S1. Figures S1-S3.

## STAR Methods

### Experimental model and study participant details

In this study, I analyzed data from the same experimental database as that described in our earlier publication^14^. Thus, no new experiments were conducted for the current report.

Briefly, the original database consisted of SC neuronal recordings from two adult, male rhesus macaque monkeys (*macaca mulatta*) performing standard visual and oculomotor tasks.

All of the original experiments were approved by ethics committees at the regional governmental offices of the city of Tübingen^14^.

## Method details

### Database details

The recordings were performed using penetration of the SC with a single, tungsten microelectrode (∼1-1.5 MOhm) per session. In the previous publication^14^, we analyzed spiking activity (firing rates) and local field potential (LFP) modulations in the stimulus-evoked epoch, to characterize visual responses in the SC. We also analyzed spiking activity in the saccade-related motor burst epoch, but only when the saccade target was visible (in the delayed, visually-guided saccade task; see below). Here, I analyzed LFP modulations in the peri-saccadic motor burst epoch; these modulations were never analyzed or documented previously. I also used the LFP results to motivate testing spiking motor bursts for saccades in the absence of a visual target (in the memory-guided saccade task; see below). Again, these analyses were not documented previously elsewhere. Finally, I analyzed LFP’s during the delay period, while the monkeys were waiting for the instruction to release their visually-guided or memory-guided saccades.

In what follows, I briefly describe the behavioral tasks relevant for this study, as well as the neurophysiological signals used. I also detail the analysis methods that I followed in the current study.

### Delayed, visually-guided and memory-guided saccade tasks

I analyzed SC spiking activity and LFP modulations around the time of saccadic eye movements generated by the monkeys during two distinct behavioral tasks.

In the first task, the saccades were visually-guided, and delayed by an explicit instruction. Specifically, the monkeys first fixated a small white spot presented over a gray background for 300-1000 ms. Then, another small white spot appeared at an eccentric location, and it remained on until the end of the trial. 500-1000 ms after the onset of the eccentric spot, the initial fixation spot was extinguished, instructing the monkeys to generate a visually-guided (albeit delayed) saccade to the (still visible) eccentric spot.

In the second task variant, the memory-guided saccade paradigm, the eccentric spot was only presented for ∼50 ms. Then, after some memory delay (300-1000 ms), the initial fixated spot was removed, instructing the monkeys to generate a memory-guided saccade towards the remembered location of the brief flash. The target reappeared only after a few hundred milliseconds of stable post-saccadic fixation. Thus, the saccade in this task variant was now generated without a visible target (towards a blank). It should also be noted here that visual stimuli were presented using a CRT display, with very short phosphor persistence time constants, and with the targets appearing over a relatively bright gray background^14^.

Therefore, it was not the case that there were residual visual signals at the saccade target location by the time that the memory-guided saccades were actually triggered.

The saccade target location was at a fixed position per session in the memory-guided saccade task, which was chosen based on the mapped response field (RF) hotspot location^14^. The latter was inferred from the delayed, visually-guided saccade task, in which we varied target location from trial to trial^12,14^. However, in the analyses of the current study, I picked only saccades towards the RF hotspot location^14^, especially because we do know that peri- saccadic LFP modulations are much weaker at SC sites far away from the movement endpoints^54^. Indeed, I used exactly the very same trials per session as those used to describe the motor bursts of the visually-guided saccades in the previous study^14^.

### Electrophysiological signals used

I analyzed firing rates and LFP modulations from all sessions in which we could observe saccade-related motor bursts. Thus, our single-electrode penetrations targeted the intermediate and deeper layers of the SC, with the great majority of our sites containing visual-motor neurons^14,17^.

The firing rates were obtained from the spike times using convolution with a Gaussian kernel having 10 ms standard deviation parameter.

The LFP’s were obtained from wide-band electrode signals that were sampled at 40 KHz and then filtered in hardware to a bandwidth of 0.7Hz-6KHz. We further filtered the signals in software as follows. First, we removed the 50, 100, and 150 Hz harmonics of the line noise using an IIR notch filter. Then, we applied a zero-phase-lag low-pass filter with 300 Hz cutoff frequency. Finally, we down-sampled the signals to 1 KHz^14^.

The motor bursts (firing rates) of the delayed, visually-guided saccades were analyzed previously^14^. Thus, I used the same sessions and same trials as in the previous analyses, but this time looking at LFP modulations. I had a total of 198 sessions containing saccade-related activity for these analyses (95 upper visual field electrode tracks and 103 lower visual field electrode tracks).

For the memory-guided saccades, their saccade-related activity was not previously described, either in terms of firing rates or LFP modulations. I thus analyzed them here, motivated by the peri-saccadic LFP results of the visually-guided saccades. The database had a total of 131 sessions possessing saccade-related activity in memory-guided saccades (49 upper visual field electrode tracks and 82 lower visual field electrode tracks). Visually-dependent saccade-related neurons^17,21–24,62^ were excluded because we were interested in genuine motor bursts at the time of memory-guided saccades.

## Quantification and statistical analysis

### Data analysis

I used Matlab (Mathworks) for all data analyses.

First, I plotted peri-saccadic LFP modulations in the delayed, visually-guided saccade task. For each electrode track, I found the interval from -200 ms to +200 ms from saccade onset. The saccade was always towards the RF hotspot location of the neuron(s) recorded in the session^14^. Then, I averaged the LFP signals across all trial repetitions of the same saccade type. In the figures, I plotted the mean peri-saccadic LFP signal around saccade onset from across all sessions of a given analysis, and I surrounded the mean traces with SEM bounds. For example, in Fig. 1D, I collected the average peri-saccadic LFP signal of each session in which the electrode was in the upper visual field representation of the SC. Then, I plotted the average of all such curves across all sessions in which the electrode track was in the upper visual field representation. This gave the blue curve in Fig. 1D. The error bars represented SEM across the sessions. A similar procedure was applied to obtain the red curve and error bars of Fig. 1D.

For some plots (like Fig. 1E in Results), I binned the electrode tracks according to the direction from horizontal represented by the encountered neurons. For example, if the SC site represented an eccentricity of 5 deg and direction from horizontal of +10 deg, then this session’s data was included in the direction bin of 0-20 deg from the horizon, in the upper visual field. This is a similar direction binning of the SC sites to what we performed earlier in the original publication with the stimulus-evoked LFP responses^14^.

To summarize the dependence of peri-saccadic LFP modulations on the visual field location, and particularly to demonstrate a step-like change as a function of direction from the horizon (e.g. Fig. 1F in Results), I measured the magnitude of the LFP signal in the interval 0-50 ms after saccade onset. In each session, I averaged the LFP values in this interval. Knowing the preferred direction of the recorded SC site of the session meant that I now had an LFP measurement (y-axis value) and a preferred direction (x-axis value) per session. Across sessions, I then binned all sessions according to their RF’s direction from the horizontal meridian (20 deg bins, in steps of 10 deg; minimum of 10 samples per angular direction bin), and I averaged the LFP magnitudes obtained from all sessions within each direction bin. This resulted in figures like Fig. 1F, with error bars denoting SEM across the sessions. The approach described here is identical to what we did earlier for visual burst epochs^14^. In other analyses, such as in Fig. S1, I performed the same analysis but this time measuring the pre-saccadic LFP value. In this case, my measurement interval was from -50 ms to 0 ms relative to saccade onset.

To test that there was a statistically significant effect of upper/lower visual field location in the above analyses, I used two approaches. First, I collected all measurements (e.g. during the interval 0 ms to 50 ms after saccade onset in Fig. 1D, F, or during the interval -50 ms to 0 ms from saccade onset in Fig. S1), and I classified them as being in either the upper or lower visual field group. Then, I applied a two-way t-test to check whether there was a statistically significant difference between the upper and lower visual field electrode track groups. I also ran a one-way ANOVA across all angular direction bins in analyses like those of Fig. 1E, to test whether direction from the horizontal indeed mattered for peri-saccadic LFP’s. I used a similar approach when analyzing spiking effects (e.g. Fig. 6E, F).

I also binned sessions according to both eccentricity and direction bins, such as in the example analyses of Fig. 2. Here, I grouped the sessions into their appropriate bins using a similar approach as above, but now taking into account both the preferred eccentricity of the recorded SC site as well as its direction from the horizontal meridian. Again, this is the same approach that we used earlier for visual response epochs^14^. The bin ranges, which are shown in the figures in Results were sometimes not having any sufficient number of sessions within them. In this case, I did not plot the corresponding bar in the bar plots (e.g. Fig. 4E for the eccentricity bin <2 deg and upper visual field). If this happened, the bin of concern was also not included in the statistical tests because there was no data available from one of the visual field groups (e.g. the upper visual field group in the example of Fig. 4E’s eccentricity bin of <2 deg). I also applied the same approach for the firing rates (e. g. Figs. 2A, 2B).

Statistically, for either the eccentricity dimension (e.g. Fig. 2C) or the direction dimension (e.g. Fig. 2D), I applied a two-way ANOVA on all sessions, after grouping them according to the two main factors of the ANOVA: (1) upper/lower visual field location; and (2) eccentricity or direction bin. I then reported the p-values for each main factor.

For the memory-guided saccades, I used similar analyses as above. To assess peri-saccadic LFP modulations despite the influence of working memory on pre-saccadic LFP values (e.g. Fig. 4A), I applied a baseline shift to the LFP’s. In particular, for each session, I obtained the average of the LFP signal during the interval from -300 ms to -200 ms relative to saccade onset. I then subtracted this baseline level from all LFP measurements in the peri-saccadic interval (-200 ms to +200 ms from saccade onset). This way, all baseline-shifted LFP curves had a value very close to zero well before saccade onset, and, therefore, well before the peri-saccadic modulations in LFP’s were expected to kick in (Fig. 4A-D).

Finally, for delay-period analyses, I used the same procedures as above. In this case, I plotted LFP curves in the interval from -300 ms to +200 ms relative to the “go” signal for the saccade (removal of the fixation spot). For summary analyses (e.g. Figs. 3E, F, S3), I used the following measurement interval: -50 ms to +25 ms relative to the time of the go signal. Thus, for each session, I averaged the LFP value during this interval, and then analyzed the data across different eccentricities or directions, and so on. The numbers of sessions, in this and other analyses, were all reported above and in the original publications^14,17^.

## Supplemental figures

**Figure S1.**
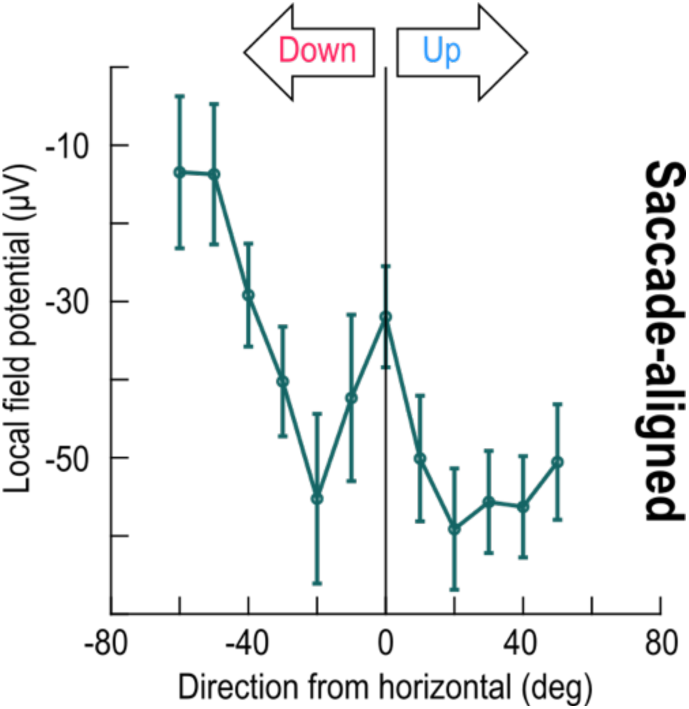
Stronger pre-saccadic local field potential (LFP) negativity in the upper visual field representation of the primate superior colliculus (SC), related to Fig. 1. This figure shows an analysis similar to that of Fig. 1F, except that I now measured the LFP in the final 50 ms before saccade onset, instead of peri-saccadically. There was still a stronger negativity in the upper visual field representation of the SC. Error bars denote SEM. Statistically, I grouped all upper visual field electrode tracks into one group and all lower visual field electrode tracks into another, and I then performed a t-test during the pre-saccadic interval (Methods). The result revealed a significant effect (p=0.0022), and that upper visual field pre-saccadic LFP negativity was ∼1.6 times larger than lower visual field LFP negativity (in absolute value). In addition, a one-way ANOVA across the shown angular direction bins revealed a significant effect of direction from the horizontal meridian (p=0.0006). Thus, the results of Fig. 1D-F also held in the pre-saccadic interval when the peri-saccadic LFP modulation was still just building up.

**Figure S2.**
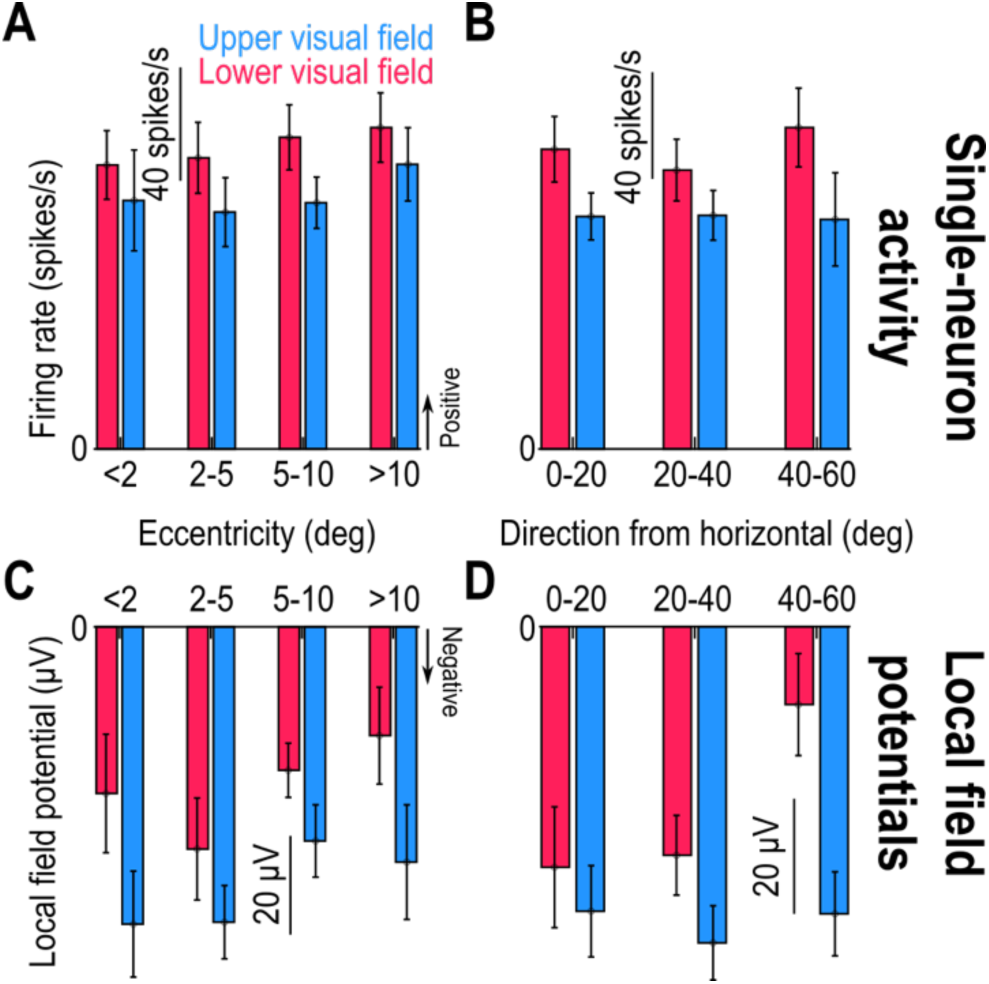
Consistency of the results of Fig. 2 in the pre-saccadic interval, related to Fig. 2. I repeated the analyses of Fig. 2, but now taking measurements during the final 50 ms before saccade onset. There were similar results to those seen in Fig. 2. Note how the LFP asymmetry was generally stronger than the spiking asymmetry (compare blue and red bars in each eccentricity or direction bin). Statistically, I compared the LFP measurements in **C** using a two-way ANOVA with the two main factors of the ANOVA being visual field location (upper or lower visual field) and eccentricity represented by the electrode track location penetrating the SC. I obtained a significant main effect for upper/lower visual field location (p=0.0086) but not for eccentricity (p=0.1892). Similarly, for **D**, I performed a two-way ANOVA with the main factors of the ANOVA being upper/lower visual field location and direction from the horizon. Again, there was a significant main effect of upper/lower visual field location (p=0.0087) but not direction (p=0.2389). For the spiking activity, there was a marginal main effect of upper/lower visual field location (p=0.0994) but not eccentricity (p=0.7266) in **A**; in **B**, there was a significant main effect of upper/lower visual field location (p=0.0119) but not direction (p=0.8579). Thus, in both the spiking and LFP’s, there were consistent asymmetries between the upper and lower visual field electrode track locations during pre-saccadic epochs. Error bars denote SEM.

**Figure S3.**
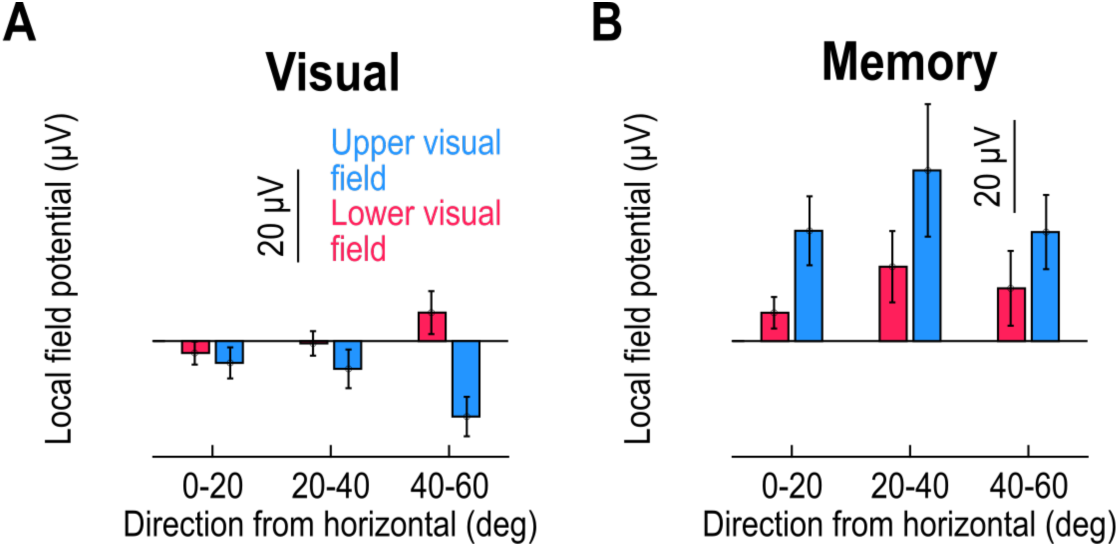
Stronger delay-period LFP negativity in the upper visual field during visual stimulation, and stronger delay-period LFP positivity in the upper visual field during working memory, related to Fig. 3. **(A)** I measured the LFP value at the end of the delay period in the visually-guided saccade task (from -50 ms to +25 ms relative to the saccade “go” signal; Methods). Across all sites, there was stronger LFP negativity in the upper, rather than lower, visual field (Fig. 3). I then binned the electrode sites according to which direction from horizontal was represented by the recorded SC activity (similar to Figs. 2, 4, 6, S2). For all upper visual field direction bins, the LFP value at the go signal was more negative than for the lower visual field direction bins. This was confirmed statistically using a two-way ANOVA (main factors being upper/lower visual field location and direction from the horizon): there was a main effect of upper/lower visual field location (p=0.0018) but not of direction bin (p=0.9038). Thus, there was stronger LFP negativity in the SC’s upper, rather than lower, visual field representation (Fig. 1A-C), even during sustained visual stimulation conditions (beyond the initial sensory-evoked transients). **(B)** In the memory-guided version of the saccade task, there was no visual stimulus driving the recorded neurons at the time of the go signal. In this case, the LFP amplitude during the delay period became positive for all electrode sites (Fig. 3), but there was still a strong upper/lower visual field effect: across direction bins, there was stronger LFP positivity for the upper visual field sites than for the lower visual field ones (p=0.0193 for the main effect of upper/lower visual field location, and p=0.3112 for the main effect of direction bin; two-way ANOVA). Thus, working memory altered LFP values in the SC (compare **A** to **B**), but there was still a clear difference between the upper and lower visual field SC representations, even in the absence of a visual stimulus. Error bars denote SEM.

